# A Pan-Cancer, Pan-Treatment (PCPT) Model For Predicting Drug Responses From Patient-Derived Xenografts With Application To Cancer Types With Sparse Training Data

**DOI:** 10.1101/2025.03.04.641355

**Authors:** Shruti Gupta, Vikash K Mohani, Ghita Ghislat, Pedro J Ballester, Shandar Ahmad

## Abstract

Predicting personalized treatment-specific responses in cancer patients requires not only robust experimental models such as PDX, which generate accurate training data on different treatment responses for the same model, but also computational models that can translate this knowledge into decisions for real patients in a timely manner. The translatability of PDX data into patient-specific outcomes is limited by the challenges in the data size requirements for training machine learning models. Previously, ML models have been developed for the two most abundant cancer types, viz. BRCA and CRC, but the same could not be scaled up to other cancer types with a smaller number of PDX samples. Here, we provide an ML framework to train a single pan-cancer, pan-treatment model for predicting treatment outcomes. We show that such models give promising results for all cancer types considered and reproduce the accuracy levels of individually trained cancer types. In the proposed model, all PDX genomic profiles from all cancer types are used as the training data, and instead of partitioning them into cancer types for each model, the cancer type and treatment name are appended as the input features of the training model. Using genomic-only and treatment-only embeddings and combining them with PCA-based dimensionality reduction, our models show promising results and provide a framework for further improvements and real-time use for best treatment selections for cancer patients.

**Data and Code Availability:** All raw and processed data, along with analysis scripts, are available at: https://github.com/Sciwhylab/pcpt

## 1. Introduction

Despite significant progress in cancer therapeutics, the prognosis remains poor and highly unpredictable (Arrowsmith, 2011; Arrowsmith & Miller, 2013; de Bono & Ashworth, 2010). It has now become clear that the cancer state and general profile of every patient determine if and which drug they will respond to effectively (Evans & Relling, 2004; Roychowdhury et al., 2011; Roychowdhury & Chinnaiyan, 2016). Some basic profiling of patients into their genetic groups and cancer types has improved clinical outcomes, but the goal of patient-specific treatment selection remains far from realized (Evans & Relling, 1999, 2004).

Accurate a priori prediction of outcomes for each possible treatment option requires experimental data that represents the specific patient’s tumor and other conditions, together with drug responses and the development of a computational model through biomarkers or whole profile methods to predict patient outcomes in real-time clinical conditions (Vargas & Harris, 2016). Traditional cell line experiments, ADMET studies, and animal pharmacological models have been used to test drug response under different scenarios. However, animal and cell-line models do not effectively represent the tumor environment or patient’s profile due to differences in their genomic, mutational, and transcriptomic profiles (Daniel et al., 2009).

The need for developing drug-response experimental models most closely representing real individual patients has led to several novel systems, of which patient-derived xenografts (PDX) are among the most powerful ones (Hidalgo et al., 2014; Imamura et al., 2015; Siolas & Hannon, 2013). PDX technology allows the transfer of cancer tissue onto a number of immunocompromised mouse, where it is grown as a patient’s experimental model. The potential effect on patients is tested by treating individual mice with corresponding treatment and recording their responses. PDX models are known to have the most remarkable similarity to human diseases yet achieved. In this way, the patient’s response to multiple treatments can be tested, which is impossible for real patients. There are many noted examples where PDX models have been used as guides for treating patients(Bertotti et al., 2011; Byrne et al., 2017; Cassidy et al., 2015; Cho et al., 2016; S. Li et al., 2013; Ma-rangoni et al., 2007; Pauli et al., 2017). PDXs maintain the genetic and epigenetic mutations and other features that contribute to drug re-sistance, and their response to drugs is correlated with those observed in the clinic(Bertotti et al., 2015; Cottu et al., 2014; DeRose et al., 2011; Ding et al., 2010; Tentler et al., 2012; ter Brugge et al., 2016).

PDX models are accurate but time-consuming and, therefore, not suitable for a real-time selection of the best treatment. However, data generated from controlled experiments on PDX can be used to learn a relationship between the genotype and transcriptome profile of tumor on the one hand and the differential response to different therapeutic options on the other hand, e.g., predicting if a patient with a given ge-nomic and transcriptomic profile would be resistant or sensitive to a specific treatment. In the past, Machine learning (ML) and Deep learning (DL) algorithms have been trained on high throughput data to develop models that can predict cancer cell lines’ response to novel drugs and drug combinations(Chiu et al., 2020). Among the many machine learning algorithms used in drug response prediction, linear regression, SVMs, ridge regression, lasso regression, elastic net, naive Bayes, neural networks, DNN, RF, GBMs, k-means, hierarchical clustering, and many more, decision trees and Gradient boosting machines(Huang et al., 2017, 2018; Jia et al., 2021; Kim et al., 2021; Y. Li et al., 2023; Sharifi-Noghabi et al., 2020; Vidyasagar, 2015) have also been used. Decision trees represent events/decisions as nodes/branches/endpoints and are easily interpretable and computationally efficient. The random forest (RF) model relies on developing an ensemble of decision trees, which has performed well on many biological problems.

An RF-based model on PDX data sets has been developed by (Nguyen et al., 2021) to predict the drug response for each treatment and cancer type pair on the NIBR-PDXE dataset (Novartis Institutes for Biomedical Research PDX Encyclopaedia) (Gao et al., 2015). The model led to the most accurate predictions when the most predictive features were used and selected through a criterion of Optimal Model Complexity (OMC), a strategy developed by the authors to build ML models using the most relevant features. They also showed that multiple gene classifiers had higher recall in vivo, consistent with in-vitro-based similar studies. They identified the most suitable profile, developed a classifier for each treatment, and assessed which molecular profile is most predictive for a given drug. One of the limitations of the models used by Nguyen et al. (Nguyen et al., 2021) was that the number of PDX experiments available for each cancer-treatment pair was small, and hence, for a large number of them, no model could be developed.

In this work, we propose a novel solution to the problem of insufficient data in developing PDX-based predictive models by introducing a framework that pools training data from all cancer types and treatments. Ordinarily, integrating multiple data sets for different classes of cancer and treatment leads to multi-class problems that do not overcome sparsely populated classes in the data because the training samples in each class remain insufficient. Our approach is to solve this problem by treating the cancer types and treatments as part of the input features instead of target classes, thereby developing a single binary class model with a larger number of training instances. We believe this approach will allow us to train models to capture interactions between similar genomic features across cancer types and intrinsically determine their contributions specific to cancer and treatment types. This so-called pan-cancer pan-treatment (PCPT) model uses a unified feature representation set by appending the cancer types and treatment types to transcriptomic and other personalized input features. To overcome the high dimensionality of the input dataset, we used PCA for feature extraction. We observed that the integrated PCPT model could be trained with promising accuracy levels across cancer and treatment types and reproduce the performance of the models with sufficient data sets reported earlier with comparable levels (Nguyen et al., 2021). Although developed for PDX data sets, the proposed approach is intuitively scalable to other multi-class problems with sparse data sets.

## 2. Methods

The overall aim of the PDX-based predictive models is to characterize a patient-specific tumor in terms of its genomic and related features. Then, for each treatment tested, it computationally predicts the outcome, whether the patient is sensitive or resistant to the corresponding treatment. Specifically, the problem to be addressed here is to take the genomic features of a PDX, including gene expression profiles, copy number variation, and mutations observed, and then given a treatment and the cancer type to which a PDX sample belongs, predict if this treatment will be effective (PDX responsive) or not (PDX non-responsive). The overall strategy of our model is two-fold. First, the high-dimensional feature data on the genomic profiling of each PDX is condensed by a PCA-based approach. Secondly, instead of training these reduced features on each cancer type and for each treatment for each PDX, a cumulative model is developed in which cancer and treatment are fed as features instead of a target class (See Fig 1). This allows a single model to be applied to any cancer type and treatment used in this cumulative model. A detailed description of the data sets, pre-processing, and training steps used are described below.

**Fig. 1.**
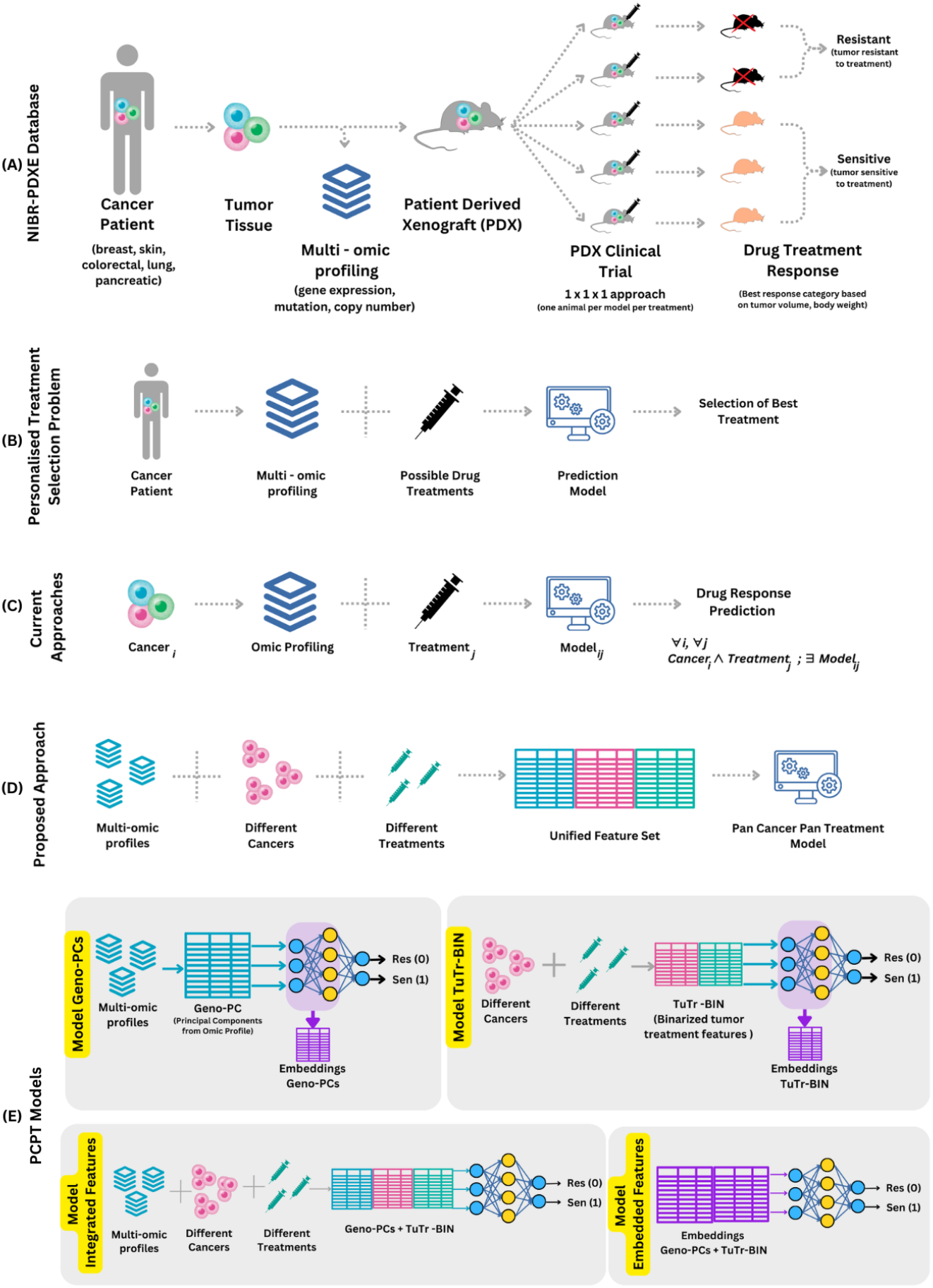
Graphical Abstract (A) Overview of NIBR-PDXE Dataset generation (B) Personalized treatment selection problem (C) Current approaches (D) Proposed PCPT approach (E) Representation of different PCPT Models.

### 2.1 Dataset Description

We used the NIBR-PDXE dataset (Novartis Institutes for Biomedical Research PDX Encyclopaedia). The dataset contains around 400 PDX models of different tumor types characterized by their mutation, copy-number alterations, and mRNA expression information and their response to drug treatments measured in terms of tumor size. The dataset studies five tumor PDX models, namely BRCA (Breast Carcinoma), CM (Cutaneous Melanoma), CRC (Colorectal Cancer), NSCLC (Non-Small Cell Lung Carcinoma), and PDAC (Pancreatic Ductal Carcinoma). The different treatments applied on these PDX are BGJ398, binimetinib, BKM120, BYL719, BYL719 + LJM716, CGM097, CLR457, HDM201, INC424, LEE011, LKA136, LLM871, paclitaxel, encorafenib, WNT974, cetuximab, CKX620, LFW527 + binimetinib, and BKM120 + binimetinib. The final dataset used in this study is summarized in Table 1, and a detailed sample count for each combination is provided in Fig 2. These statistics are derived from the original data available as Supplementary Table in xlsx format containing five sheets named RNASeq_fpkm, copy_number, pdxe_mut_and_cn2, PCT_raw_data, and PCT_curve_metrics which contains gene expression data, actual gene copy number, mutations, and categorical copy number alterations, raw response in terms of tumor volume change, and the processed response of PDX into different classes, respectively in the previous works, where additional details of data sets are available (Gao et al., 2015, Nguyen et al. 2021). This data is referred to as Gao data in the rest of this work.

**Table 1.**
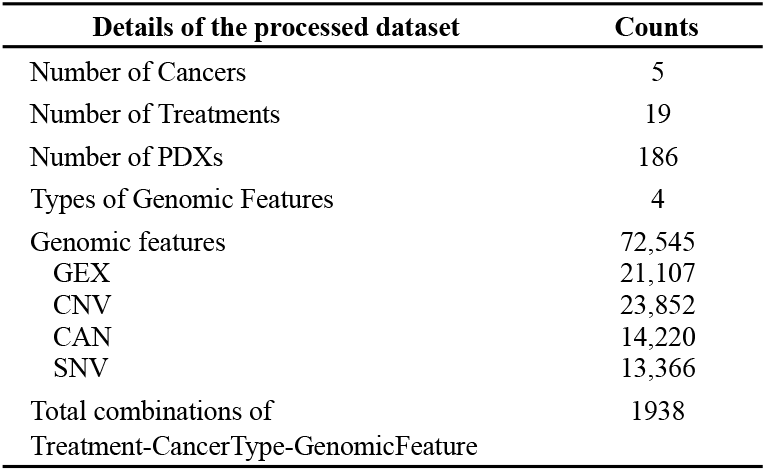
Summary of the experimental data used in this study.

**Fig. 2.**
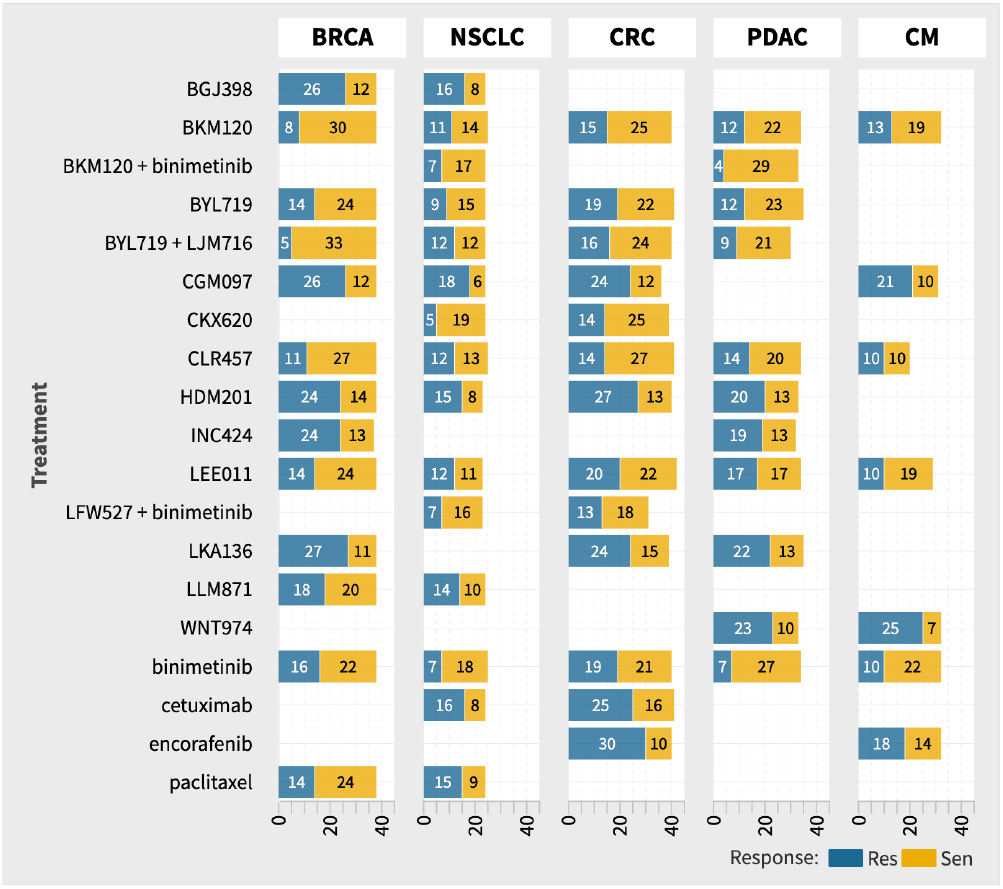
Number of samples tested for each treatment-cancer pair grouped by their response to treatment

### 2.2 Data Pre-processing

Gao dataset generated five different genomic information matrices to describe individual tumors, namely Gene Expression Matrix (GEX), Copy Number Variation (CNV), Copy Number Alterations (CNA), Single Nucleotide Variation (SNV), and PDX Clinical Trial (PCT) (Gao et al., 2015, Nguyen et al. 2021) (see Table 1). They were pre-processed or used as such as follows:

#### Gene Expression Matrix (GEX)

The GEX matrix in the RNASeq_fpkm sheet contains normalized gene expression values in FPKM (Fragments Per Kilobase of transcript per million mapped reads). It contained expression data for 22,665 genes across 399 PDX samples and was used without additional pre-processing.

#### Copy Number Variation (CNV)

The CNV matrix in the copy_number sheet contains the copy number of 23,852 genes across 375 PDXs. The data were directly used as provided in the sheet.

#### Copy Number Alterations (CNA)

The CNA matrix was derived from the pdxe_mut_and_cn2 sheet, where categorical copy number data (e.g., Amp5, Amp8, Del0.8) were binarized. Aberrant copy numbers were marked as 1, and normal values as 0. The final matrix contains 21,818 features for 399 PDXs. The data were directly used as provided in the sheet.

#### Single Nucleotide Variation (SNV)

The SNV matrix is derived from the pdxe_mut_and_cn2 sheet and was generated by binarizing the mutation data. A value of 1 indicates the presence of at least one somatic mutation in a gene, while 0 indicates no mutation. The final matrix contains 21,818 binary features for 399 PDXs.

#### PDX Clinical Trial Data (PCT)

The PCT matrix is extracted from the PCT_raw_data and PCT_curve_metrics sheets and includes recalculated response categories based on Nguyen et al. (2021) namely Sen (Sensitive or Respondent) and Res (Resistant or Non-Respondent).

After combining the genomic feature matrices and merging them with the treatment profiles for each PDX model, we removed columns with missing data. This resulted in 186 unique PDX models and 1,938 tumor-treatment pair combinations. The final dataset consists of 1,938 data entries, representing combinations of treatment, cancer type, and genomic features across five cancer types. Only PDX models with complete molecular profiling and response data for at least three samples in the respondent or non-respondent categories were retained.

### 2.3 Feature Engineering

#### 2.3.1 Genomic Feature Extraction using PCA

We applied Principal Component Analysis (PCA) using the PCA module from the scikit-learn library to reduce the dimensionality of the genomic data, which initially consisted of 72,545 features (see Table 1). We selected the top 200 principal components (PCs) as representative of all these features (see Results section). Before performing PCA, the genomic features were standardized in Python by subtracting the mean and scaling to unit variance using the StandardScaler function from the scikit-learn library (Pedregosa et al., 2011). The resulting set of 200 principal components, referred to as “Geno-PC” throughout the article, was used as the input feature set for subsequent analyses.

#### 2.3.2 Tumor and Treatment Class Features

To develop a pan-cancer, pan-treatment (PCPT) model, we included categorical features representing both tumor types and treatment types. These were one-hot encoded, resulting in 24 binary features: 19 for treatment types and 5 for tumor types. This set of 24 binary features is referred to as “TuTr-BIN” in the article and is appended as an input vector (resulting in 224 features for each sample) for our classifier model.

### 2.4 PCPT model and training strategy

As outlined above, we adopted a novel strategy wherein we treated cancer and treatment types as input features and developed a unified feature representation set. This contrasts with previous studies, where a different model is trained for each combination. The predictive model was developed on PCPT representation to classify patients into respondent and non-respondent classes.

We used a simple feed-forward neural network consisting of one input layer, a hidden layer with eight nodes, and an output layer. In our study, we used either the rectified linear unit (ReLU) or the tanh activation function in the hidden layer and applied a sigmoid activation function in the output layer to train a binary classification model.

To optimize the network’s trainable parameters (weights and biases), we used the binary cross-entropy loss function to measure the difference between predicted probabilities and true binary labels. We applied a dropout rate of 0.2 in the hidden layer to prevent overfitting. Dropout helps by randomly deactivating neurons during training, effectively averaging multiple models and reducing overfitting (Srivastava et al., 2014). We initialized the weights using Glorot normal distribution sampling (Glorot & Bengio, 2010). We trained the model using the Adam optimizer with a fixed learning rate of 1.00E-04. The network was trained for 500 epochs with a batch size of 64.

We explored different feature representation strategies of the features and trained models for each representation. These models are described in Table 2. Unless otherwise noted, all models were trained with the same parameters. This structured approach allowed us to test and compare various feature sets, leading to the development of the final PCPT model that combines predictions from the previous models for enhanced performance. Apart from the original 200-PCA features and 224 features, which included the tumor and treatment-derived features, additional approaches based on auto-encoder-like embeddings were also used, as listed in Table 2 and described below.

**Table 2.** Summary of novel models, feature representation strategies, and hyper-parameters. Number of hidden layers in each model is one with eight nodes. Activation and loss function in each model are sigmoid and binary cross-entropy.

| Model | Input Feature set description | Input size<br>(features x samples) | Activation<br>function |
| --- | --- | --- | --- |
| Model-Geno-PC | Geno-PC feature set | 200 x 1938 | ReLU |
| Model-TuTr-BIN | TuTr-BIN feature set | 24 x 1938 | tanh |
| Model-Integrated-features | Combined Geno-PC and TuTr-BIN feature sets | 224 x 1938 | ReLU |
| Model-Embedded-feature | Embeddings from Geno-PC and TuTr-Bin models | 16 x 1938 | tanh |
| Model-PCPT | Average of the prediction scores of the above four models | 4 x 1938 | - |

### 2.5 Embedding Extraction Process

In our model development, we created a Model-Embedded-Feature by extracting embeddings from the Model-Geno-PC and Model-TuTr-BIN, the response class predictions trained exclusively from genomic PCA (200 features) and tumour-treatment (24 features), respectively. The embeddings represent compressed, lower-dimensional representations of the original feature sets, allowing us to efficiently integrate the information from genomic features and tumor-treatment classifications.

For the Model-Geno-PC, we trained a neural network on the genomic principal components (Geno-PC), and the activations from the hidden layer were used as the embeddings. Similarly, for the Model-TuTr-BIN, the activations from the hidden layer were extracted as embeddings representing tumor and treatment-type features(TuTr-BIN).

These embeddings were then concatenated, resulting in a combined embedding of size 16 for each sample. This embedded representation was used as input for the Model-Embedded-Feature, where the compressed information from genomic and class features, together with cross-validation-based training, allowed for efficient classification.

### 2.6 Model training and cross-validation

We employed 4-fold cross-validation to evaluate model performance, using a proportional sampling approach for tumor-treatment pairs. Samples from each tumor-treatment pair were distributed proportionally across the folds. Pairs with fewer samples contributed fewer instances per fold, while pairs with more samples contributed more. This ensured that each fold maintained a proportional representation of the tumor-treatment pairs, preventing any pair from being over- or under-represented during training and evaluation.

During each fold training and testing, we maintained a strict separation between training and test data, preventing information leakage and ensuring accurate model performance estimates e.g. by computing embeddings for each iteration only from the models based on training data, ensuring the integrity of the evaluation process and helping us to estimate true performance levels on unseen samples.

### 2.7 Evaluation of the performance of different PCPT models

We evaluated the models using key metrics, including the F1 Score and Matthews Correlation Coefficient (MCC), chosen for their robustness in handling imbalanced classes. The decision threshold was optimized based on the true positive rate (TPR) and false positive rate (FPR). Performance was assessed globally across all tumor-treatment pairs, individual tumor types, and treatments to identify any model biases and ensure broad generalizability.

## 3 Results

As described in the Methods, we have tested multiple methods of creating an integrative prediction to arrive at treatment response status from genomic feature sets (Table 2). In each model, our starting representation of genomic features is the cross-validated principle components (transformation matrices derived only from the training data). In the following section, we systematically describe the results and analysis of the training models.

### 3.1 Principal components of Genomic features of PDX’s

We extracted principal components from 72,545 features covering the four molecular profiles of the samples. First we examine the entire data to explore the true dimensionality of genomic features. Fig 3(a) shows the scree plot representing the cumulative variance captured by increasing the number of selected components. A total of 10, 20, 50, 100, and 150 principal components explain 23.23%, 36.83%, 62.51%, 84.94%, and 96.56%, respectively. The top 200 components cover most of the variance in the data and form the feature representation of our genomic information for subsequent predictive models.

**Fig. 3.**
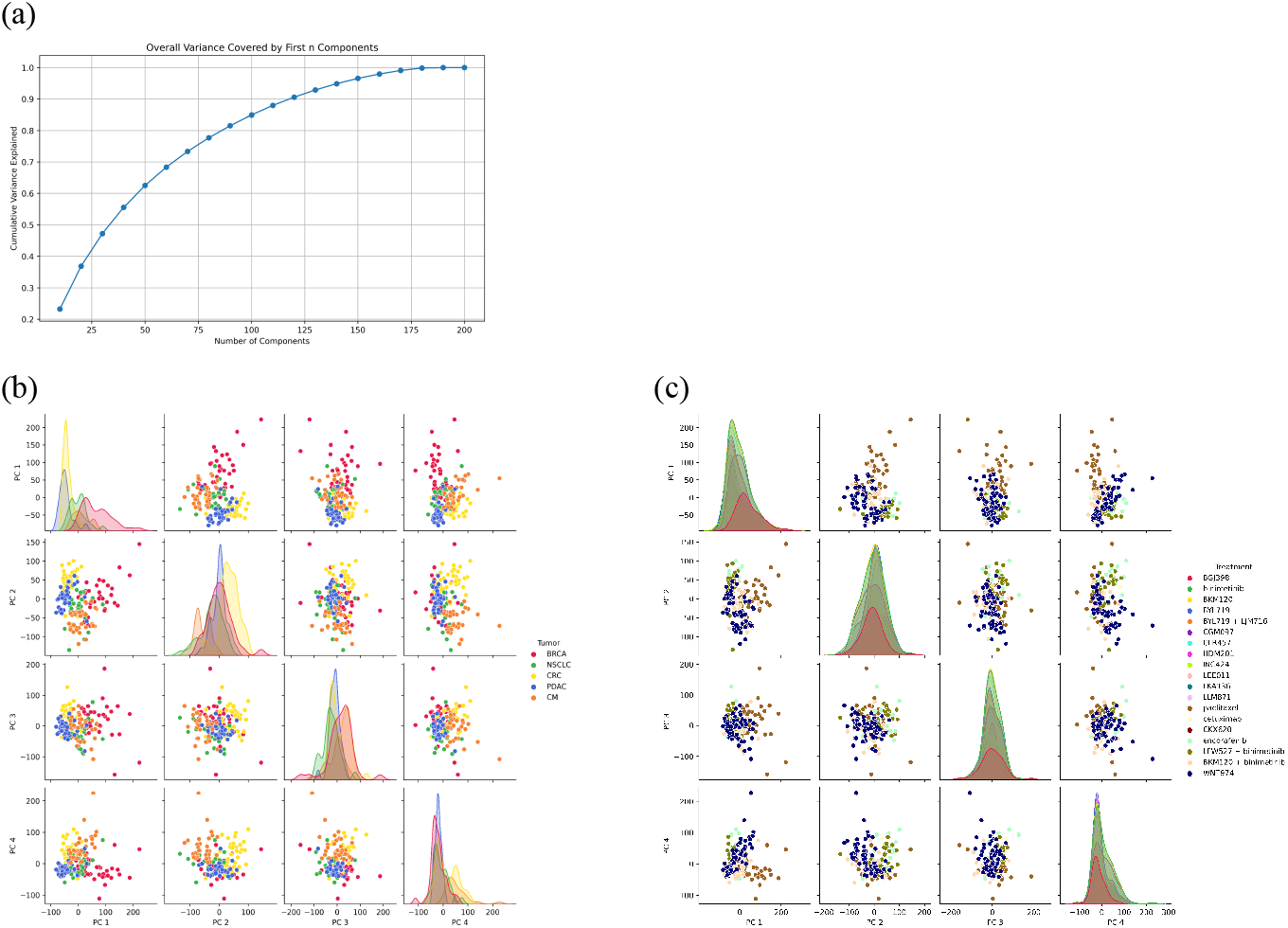
Principal component analysis of genomic features in all PDX samples (a) variance explained by different number of top principal components (b) separation of different cancer types in top 4 components of PCA space (c) distribution of samples given different treatments across all cancer types in the top 4 components of PCA space

The principal component of genomic features by pooling the all-cancer types and treatments presents before us the opportunity to explore cancer-specific variations in PC space. These individual variations are critical to differential responses to each treatment and transfer of training across cancer types right at the feature representation level. Different tumor types are expected to have some genomic profiles specific to cancer type, irrespective of their response to treatment.

However, treatments are an external intervention and, therefore, are blind to genomic PCs. Any distinction in PCs in treatment space may simply mean the differences in the choice of patients made for giving that treatment. They may also represent random variations in genomic profiles captured as biases due to a small number of samples in some treatments. In Fig 3(b), we observe a clear, narrow peak in density plots of PC1 and PC2 for CRC and PDAC cancers, respectively, which suggests less variability within the cancer type concerning the molecular features captured by that component. In addition, we observe that PC1 can well separate the BRCA cancer types, and PC4 also shows a clear, narrow peak for BRCA cancer. A separation of CRC and CM cancer in PC2-PC4 is also observed. Our treatment response model aims to distinguish between tumor responses and not the tumor itself, although the intrinsic differences in tumor profiles are inevitable. This analysis shows that the treatment-independent separation of cancer types within the principal components is a major challenge that any predictive model must overcome in effectively learning a treatment-specific response across cancer types. Fig 3(c) confirms that the treatments are generally not much separated between the top principal components and, therefore, meets an important prerequisite for the data quality and consistency of the proposed models. However, we observe that the paclitaxel drug is somewhat separated from the rest of the clusters in the scatter plot, although there is no distinct peak in density distributions. This suggests that the nature of patients given this treatment was slightly biased, irrespective of their response levels to treatments. This bias is observed only in one treatment; therefore, we have made no attempt to correct it in the current work.

### 3.2 Prediction Performance of the best PCPT model

As shown in Table 2, we have used five different types of PCPT models. We discuss the outcomes of each of these models in a later section. We first discuss the performance levels of the best PCPT model (simply called the PCPT model) in our work, and it is based on the predicted scores for each PDX sample obtained from four different trained models. It may be noted that all prediction scores are based on the predictions for the samples when they were in the test data sets and represent the true generalizability of the model under a cross-validation scheme, preventing information leakage from the training steps. Fig 4(a) shows that the final PCPT model performance is somewhat varied across tumor-treatment pairs. While the F1 scores, accuracy, and AUC indicate a reasonable level of performance, the low MCC values could be due to class imbalances. It should be noted that for highly class-imbalanced data, MCC scores often underestimate predictive power, and thus, the other scores must be considered as the guide (Dana et al. NAR GAB, 2022). Fig 4(b-f) shows the ROC of the PCPT model for BRCA=0.62, CM=0.62, CRC=0.70, NSCLC=0.60, and PDAC=0.63. Prima facie, a performance range between 0.62 and 0.70 observed here may look modest, but these results must be seen in the following context. First of all, for some of the tumor-treatment types, the number of samples is too small, and it has so far been not possible to train a model exclusively on such data. Thus, whatever prediction performance we get here is good because these models have been trained using ML for the first time and, as of now, the only way tried. Secondly, in a high dimensional system such as this, over-fitting is a great risk, and high-performance levels are often misleading because generalization is low. Current models have been trained with strict cross-validation controls and represent a high confidence of generalization. Third, the genomic profiles are not the only factor that determines the effectiveness of a treatment, as there are many confounding factors, such as non-genomic factors, individual host variation, and cellular heterogeneity. In the current model, genomic features alone have been shown to provide us with useful leads to treatment selection, which can be combined with other clinical leads to provide a more powerful personal therapy. Finally, the field of PDX-derived ML models for treatment response is still in an early stage, and future models will improve the performance. In contrast, the proposed model provides a framework for working with a large number of classes with a low number of samples in each group.

**Fig. 4.**
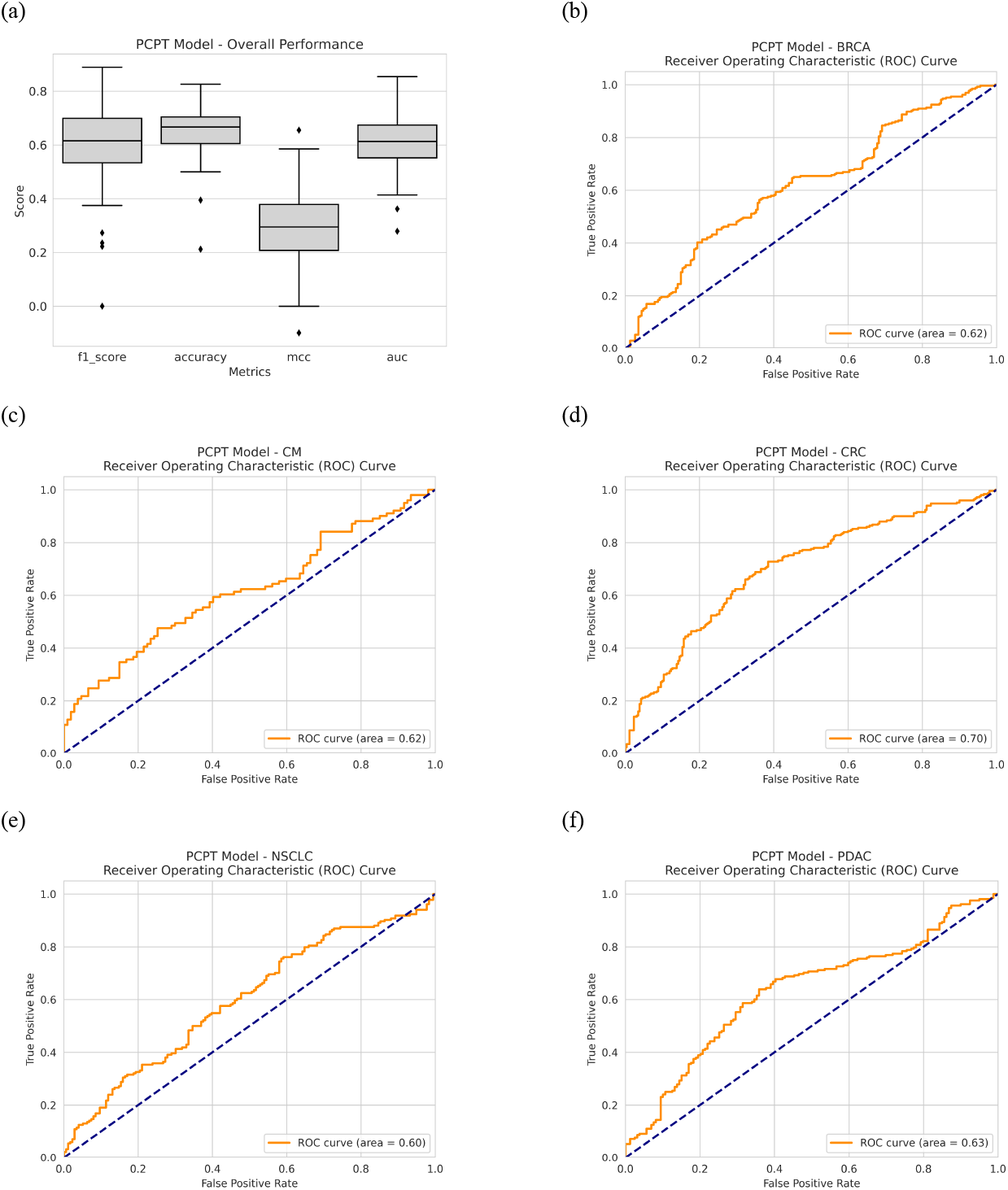
Predictive performance of PCPT model (a) overall performance scores for each tumor-treatment pair measured by different metric (b-f) AUC of ROC for each tumor type.

As stated above, many of the tumor-treatment pairs are trained here for the first time and cannot be compared with anything in the literature. However, BRCA and CRC systems have sufficient data, and effective models have been developed for them previously. We compare in the following if the PCPT model trained in combination with all cancers can retain the performance on samples of these two cancers reported by the previous RF-OMC model proposed by Lin et al. We also compare the performance levels of PCPT precursor models (first four rows of Table 2) with the finally aggregated PCPT model for these and other cancer types in the subsequent sections. Fig 5 shows the comparison of the PCPT model with the RF-OMC model of Lin et al. on BRCA and CRC tumors. Since only the F1-score and MCC have been reported in the previous work, the comparison is made only for these scores. We observe that for some treatments, PCPT outperformed the RF-OMC model and vice versa in other cases and that the prediction performance levels from the two models are comparable (See Table 3). Specifically, BRCA results are better for RF-OMC models, whereas PCPT outperforms for CRC, making it overall a negligible difference. In the case of BRCA, an overall drop in performance is actually driven by three treatments, most prominently paclitaxel (see Fig 5), which was also flagged for its special behavior in the PCA in the section above. It appears that the genomic profile data belonging to some treatments has a bias that is difficult to capture in a larger model relying on global deterministic signatures of treatment response. An apparent anomaly in the PDX data for some of the treatments, especially paclitaxel, suggested here is a matter of further investigation in the future. Here, we conclude that the overall prediction performance levels suggest that even for the cancer types with sufficient data, PCPT could reproduce at least the same accuracy levels obtained by specialized models trained on only one cancer type. This result may be used to gain further confidence in the prediction performance of cancer types never trained using ML models so far.

**Table 3.**
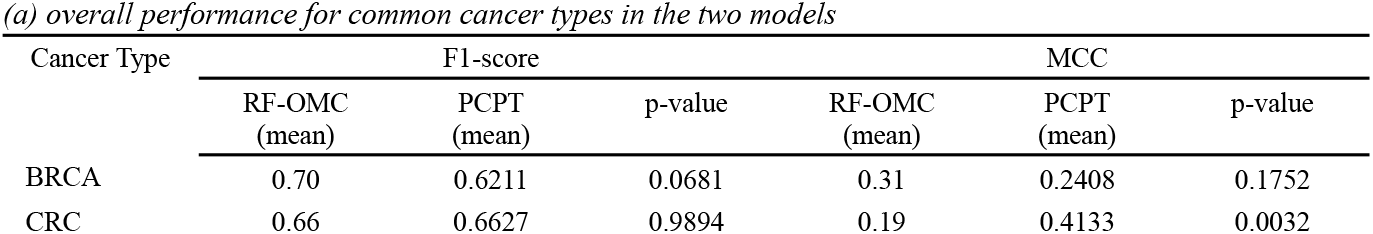
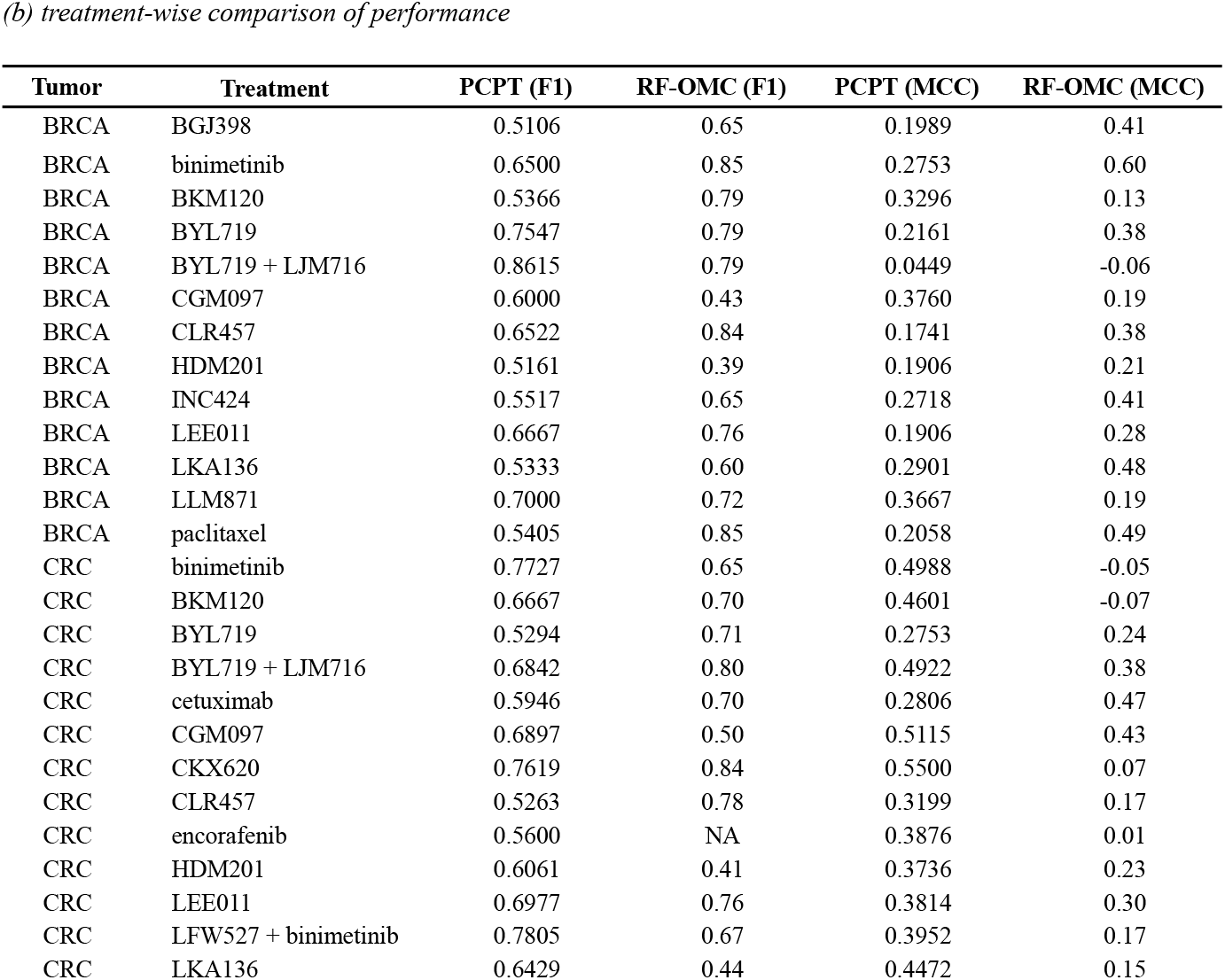
Comparison of prediction performance between PCPT and RF-OMC models for the cancer types trained previously as individual models (BRCA and CRC)

**Fig. 5.**
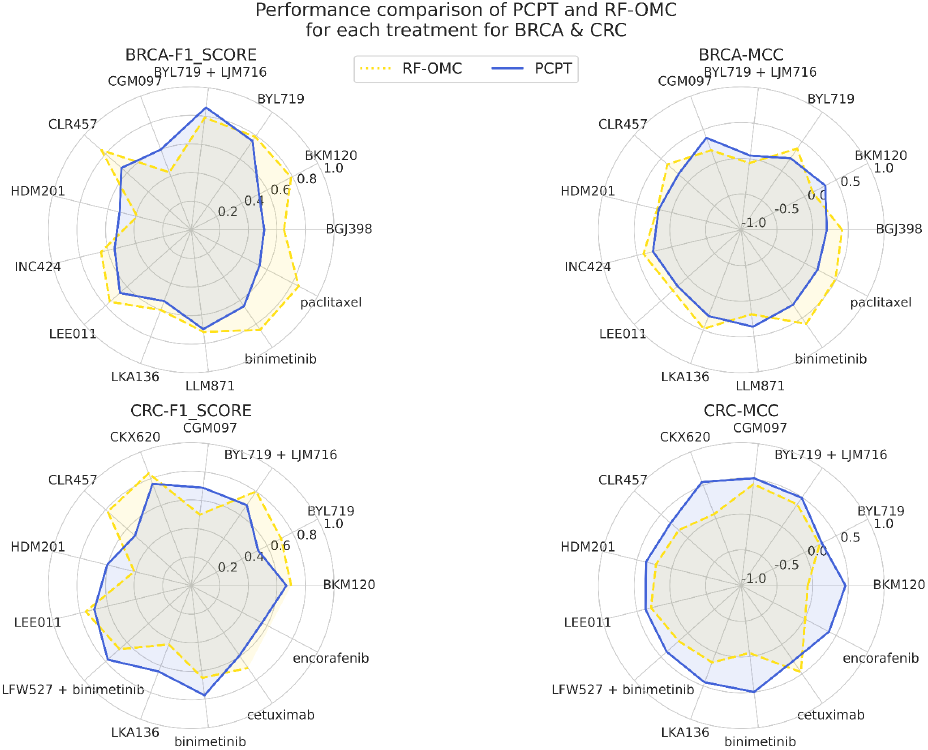
Comparison of PCPT model with RF-OMC model for BRCA1 and CRC cancers for all treatments.

### 3.3 Performance Comparison of different PCPT models

As described in Methods, we used four different predictive models by representing genomic information of PDX samples in different ways, and the final PCPT model averages the predictions obtained from each of the four models. Here, we examine the predictive power of each of the four constituent precursor models to estimate the nature of our models’ predictive information. Figure 6 shows the treatment-wise distribution of prediction performance of different PCPT models in terms of four performance metrics. We observe that the models based on the PCA representation of genomic profiles and their corresponding low-dimensional embeddings serve well and contribute maximally to the final PCPT models. Treatment alone is not predictive, which indicates that the model has effectively eliminated baseline effectiveness levels of drugs. Nonetheless, the final PCPT model is slightly better than all the precursor models and more robust as the interquartile ranges are somewhat lower than the genomics-only features.

**Fig. 6.**
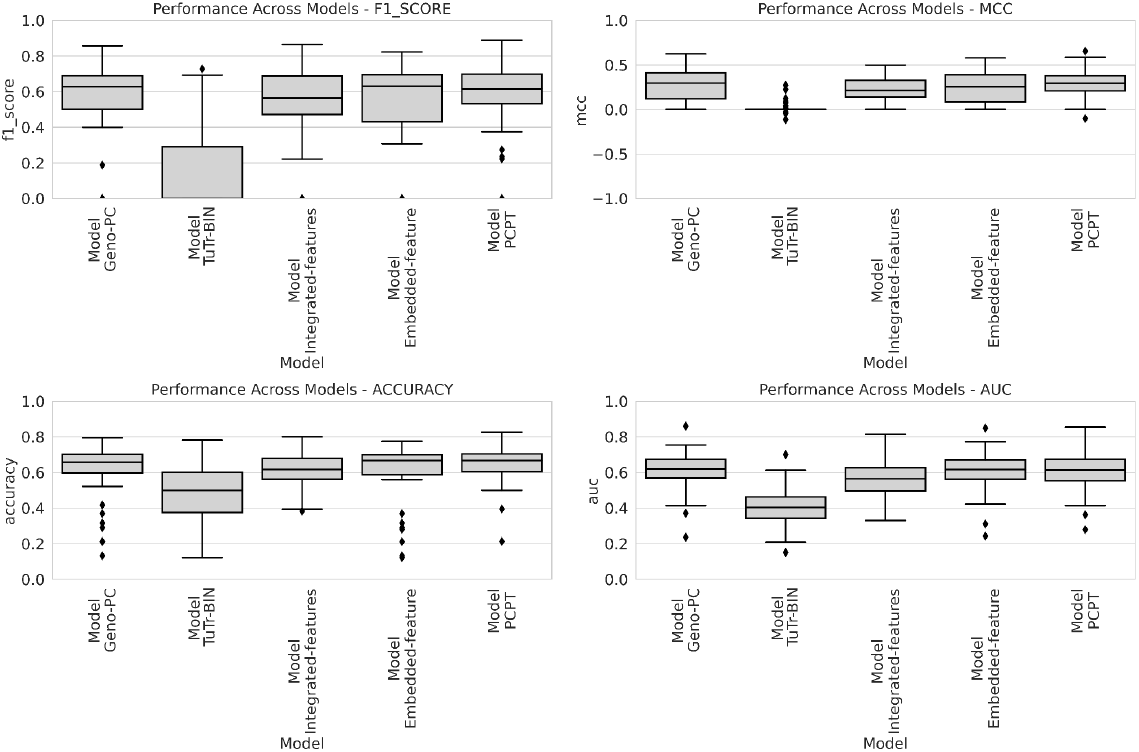
Overall performance of different PCPT models for all tumour-treatment combinations

### 3.4 Class imbalance and prediction performance

We observe that the number of sensitive and resistant samples for each treatment and cancer type is not proportionate in the data sets (see Fig 2). This class imbalance may not only impact the trainability of models but also produce misleading prediction scores, e.g., MCC has been reported to underestimate performance for class-imbalanced data sets. In Fig. 7, we show the model F1 and MCC scores as a function of relative number of sensitive data for the corresponding tumor-treatment pair. An overall look at these plots suggests that only the NSCLC models show a significant correlation between the F1 score and the number of samples in the sensitive class. Observations made from the five cancer types and only in the F1 score is difficult to assess a general trend, and we broadly conclude that the performance scores reported in this work hold true for all the class imbalance ranges covered for the data sets used.

**Fig. 7.**
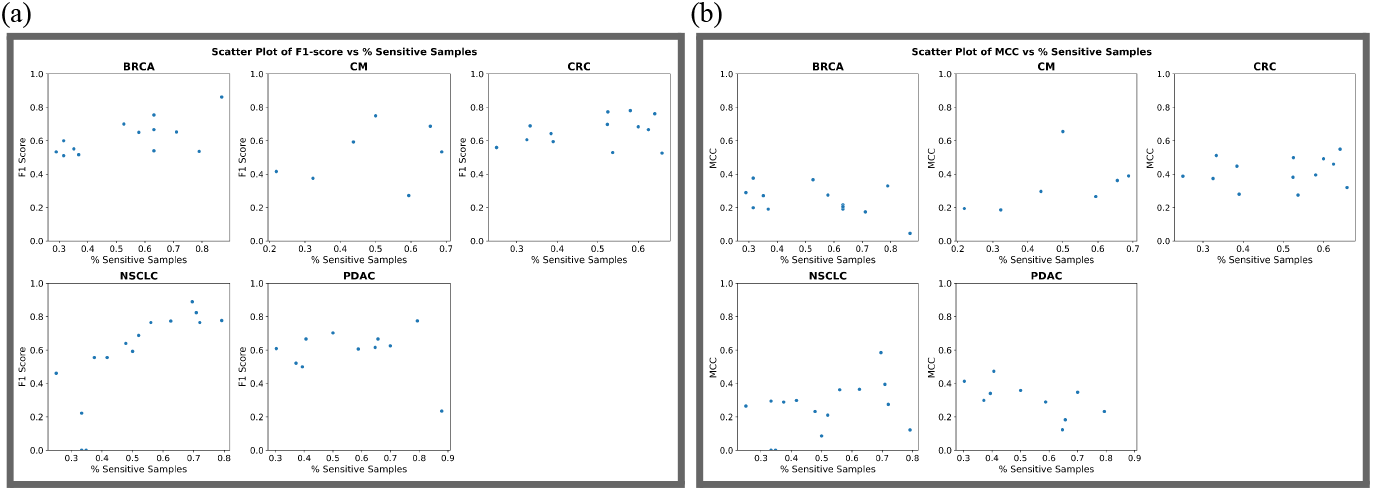
Cancer-wise trends on prediction performance trends with respect to the size of treatment data in the sensitive group (class imbalance)

## 4 Conclusion

We have successfully developed an integrative genomics-transcriptome-based approach to predict treatment responses for multiple cancer types using PDX data. Even though performance levels are modest, they are not only comparable with the state-of-the-art models available for single cancer types but also provide a much larger coverage of patient groups. Results also suggest that the genomic profiles contain significant information about the specific responses to treatments by individual patients, of which we have been able to capture at least some. Combined with other clinical contexts, these results will be helpful in the personalized selection of cancer therapies with the aim of better treatment outcomes.

## Acknowledgements

We gratefully acknowledge funding from the Indo-French Centre for the Promotion of Advanced Research (CEFIPRA) to SA and PJB and resources provided by the DBT-Bioinformatics Center at SCIS, JNU (SA, SG, VKM). SG gratefully acknowledges ICMR for Junior Research Fellowship (No. 3/l/3/JRF-2018/HRD - 031).

